# Estimating the effects of variation in viremia on mosquito susceptibility, infectiousness, and *R_0_* of Zika in *Aedes aegypti*

**DOI:** 10.1101/221572

**Authors:** Blanka Tesla, Leah R. Demakovsky, Hannah S. Packiam, Erin A. Mordecai, Américo D. Rodríguez, Matthew H. Bonds, Melinda A. Brindley, Courtney C. Murdock

## Abstract

Zika virus (ZIKV) is an arbovirus primarily transmitted by *Aedes* mosquitoes. Like most viral infections, ZIKV viremia varies over several orders of magnitude, with unknown consequences for transmission. To determine the effect of viral concentration on ZIKV transmission risk, we exposed field-derived *Ae. aegypti* mosquitoes to four doses (10^3^, 10^4^, 10^5^, 10^6^ PFU/mL) representative of potential variation in the field. We demonstrate that increasing ZIKV dose in the blood-meal significantly increases the probability of mosquitoes becoming infected and infectious, as well as the rate at which virus spreads to the saliva, but found no effect on dissemination efficiency or mosquito mortality. We also demonstrate that determining infection using RT-qPCR approaches rather than plaque assays potentially over-estimates key pathogen parameters, including the time at which mosquitoes become infectious and viral burden. Finally, using these data to parameterize an *R_0_* model, we demonstrate that variation in viremia substantially affects transmission risk.

## Introduction

Although discovered in 1947 (1), Zika virus (ZIKV) has recently become a public health concern due to its rapid spread and newly identified teratogenic effects. Shortly after isolation from a rhesus macaque in Uganda, the virus caused several mild infections in humans (2, 3). ZIKV infections remained unapparent until the first major outbreak in 2007 on the island of Yap (4). The virus further spread across the Pacific, where it was first associated with Guillain-Barré syndrome during the 2014 French Polynesian outbreak (5). In 2015, transmission was confirmed in Brazil (6), after which the virus spread rapidly across the Americas (7). ZIKV was declared a “public health emergency of international concern” by WHO in 2016 due to excessive spread and increases in complications associated with congenital Zika virus syndrome (8).

The primary route of ZIKV transmission is through the bite of *Aedes* mosquitoes. The principal urban vector in the Americas is *Ae. aegypti* while *Ae. albopictus* is believed to be a secondary vector (9). Although most cases of ZIKV infection are asymptomatic (4), 20% of individuals develop symptoms associated with Zika fever (10). Currently, human viremia is not well characterized. Studies suggest that ZIKV viremia in the blood is lower than other arboviruses and does not significantly differ between symptomatic and asymptomatic patients (11). In arboviral systems such as dengue, variation in viremia across infectious human hosts influence the number of mosquitoes that become infectious (12), yet this has only been minimally explored in the ZIKV system (13–15). Further, the impact of variation in host viremia on overall transmission has yet to be adequately addressed.

The number of people at risk for contracting ZIKV or other similarly transmitted arboviruses (e.g. dengue and chikungunya) cannot be accurately estimated, as most ZIKV infected hosts are asymptomatic, the distribution of hosts with varying viremia is unknown, and the relationship between variation in host viremia and transmission to local mosquito populations is unclear. *R_0_* (the basic reproductive number of a pathogen) represents the expected number of secondary cases that result from a single infection in a susceptible population and is comprised of a combination of human, mosquito, and pathogen traits (16, 17). *R_0_* models allow for the estimation of the epidemic spread of pathogens (16–18), are commonly used to assess the effectiveness of mosquito control strategies (19–22), and are routinely used to predict the coverage required for successful vaccination programs (23–25). Yet, our current ability to estimate the number of human hosts at risk or to control ZIKV transmission is a major challenge due to this lack of basic information on transmission mechanisms and inability to build scientifically sound mechanistic models, the most fundamental of which is R_0_. To address this limitation, we conducted experiments to assess the effect of variation in viral dose on vector competence, the extrinsic incubation rate (EIR), and the daily probability of mosquito survival. We used these results to parameterize a mechanistic *R_0_* model and to estimate the number of infectious mosquito bites contributed by mosquito populations feeding on hosts of varying viremias.

## Results

### The Effect of Viral Dose on Vector Competence and EIR

To investigate how variation in ZIKV dose affects vector competence, transmission efficiency, and the extrinsic incubation period in *Ae. aegypti*, we orally infected mosquitoes with four different viral concentrations (10^3^, 10^4^, 10^5^ and 10^6^ PFU/mL) reflecting viremia in ZIKV-infected humans. We found that the mean proportion of infected mosquitoes, mosquitoes with disseminated infections, and infectious mosquitoes significantly increased with increasing viral dose (Table 1, Fig 1). The infectious dose required to infect 50% of the mosquito population (ID_50_) was 10^4.98^ PFU/mL. We also observed a significant effect of dpi on the probability that mosquitoes had disseminated infection or became infectious, but not on the probability that they became infected (Table 1). At the highest doses (10^5^ and 10^6^ PFU/mL), the virus was detectable in mosquito bodies at all tested time points (Fig 2A). On average, more than 4 days are required for ZIKV to disseminate into the head (Fig 2B) and more than 8 days to be present in the saliva (Fig 2C). Finally, the significant interaction between ZIKV dose and dpi indicates that increases in viral concentration significantly increased the rate at which mosquitoes disseminate infection and become infectious (Table 1, Fig 2). These results suggest that mosquitoes feeding on human hosts with varying levels of circulating virus could experience both different probabilities of infection and overall infection dynamics.

**Table 1.** The effect of dose, day, and potential interaction on mosquito infection, dissemination, infectiousness, and transmission efficiency.

| response variables | dose |  |  | day |  |  | dose x day |  |  |
| --- | --- | --- | --- | --- | --- | --- | --- | --- | --- |
|  | F | d.f. | p-value | F | d.f. | p-value | F | d.f. | p-value |
| probability of infection | 33.898 | 3 | **<0.0001** | 0.004 | 4 | 1 | 0.501 | 12 | 0.9 |
| probability of dissemination | 52.61 | 3 | **<0.0001** | 11.929 | 4 | **<0.0001** | 4.295 | 12 | **<0.0001** |
| probability of infectiousness | 22.86 | 3 | **<0.0001** | 7.82 | 4 | **<0.0001** | 3.45 | 12 | **0.002** |
| efficiency of dissemination | 0.022 | 3 | 0.996 | 0.097 | 4 | 0.982 | 0.134 | 8 | 0.997 |
| efficiency of salivary gland infection | 2.566 | 2 | 0.108 | 3.136 | 4 | **0.044** | 0.558 | 4 | 0.696 |

**Fig 1.**
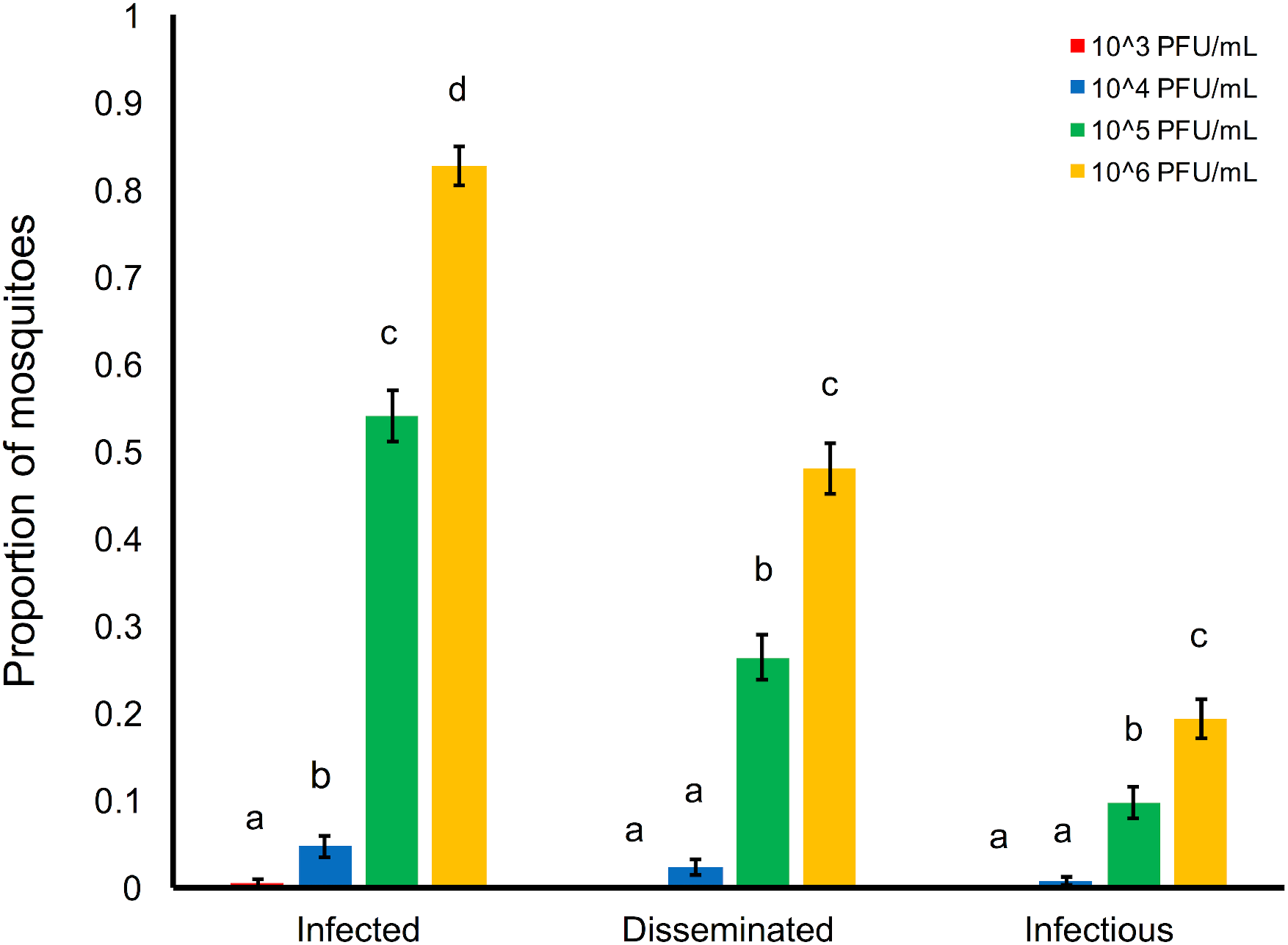
ZIKV dose and the proportion of mosquitoes infected, with disseminated infections, and infectious. Relationship between the ZIKV dose (10^3^, 10^4^, 10^5^, and 10^6^ PFU/mL) and the proportion of mosquitoes infected (ZIKV positive bodies compared to total number of exposed), with disseminated infections (ZIKV positive heads compared to total number exposed), and infectious (ZIKV positive saliva compared to total number exposed). For each category, results with no common letters were significantly different (*p* ≤ 0.05).

**Fig 2.**
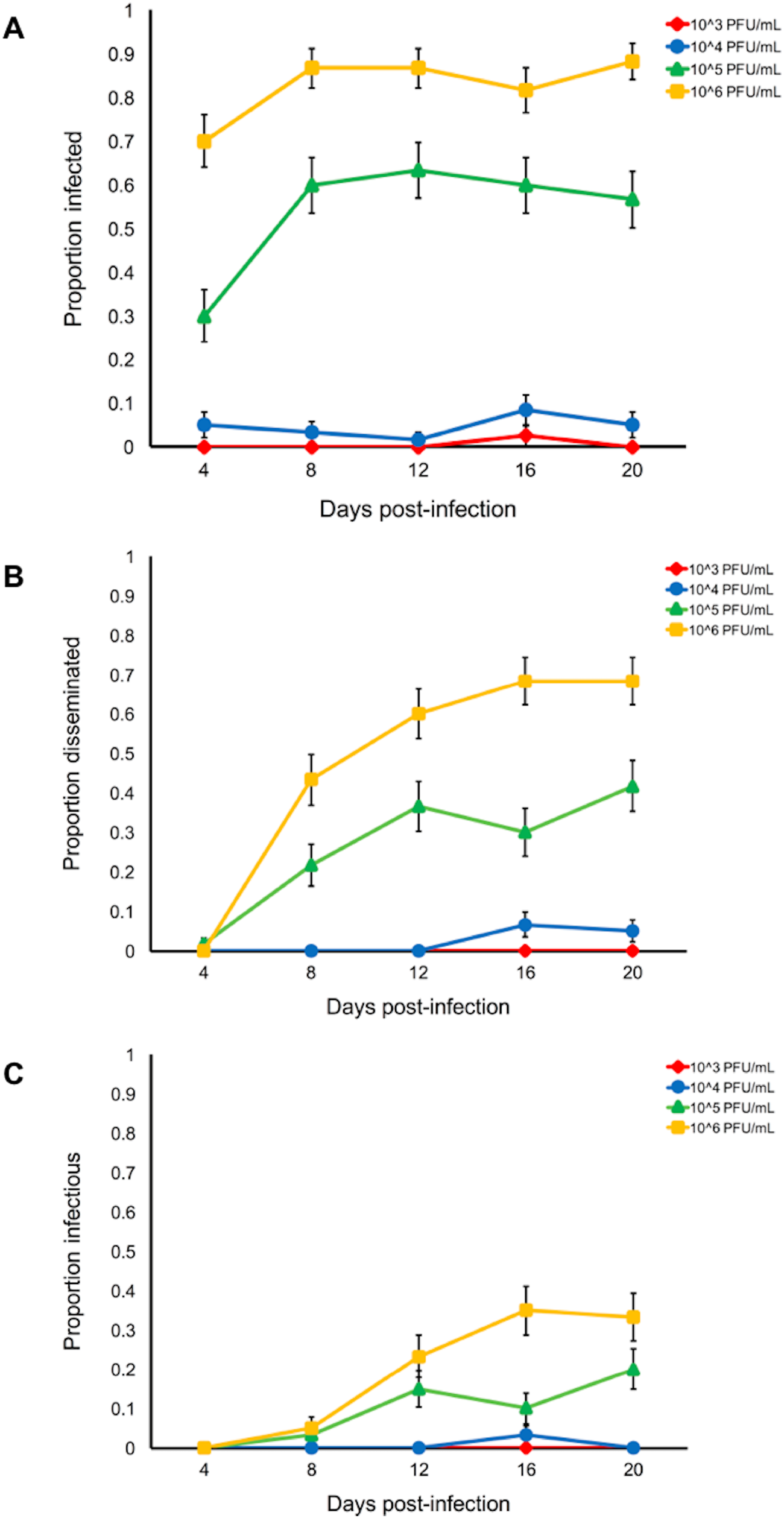
Days post-infection and the proportion of mosquitoes infected, with disseminated infections, and infectious. The relationship between days post-infection (4, 8, 12, 16, 20) and the proportion of mosquitoes infected (A), with disseminated infections (B), and infectious (C) after exposure to four different viral doses (10^3^, 10^4^, 10^5^, and 10^6^ PFU/mL).

Results from generalized linear mixed effects models examining the effects of dose, day, and the interaction on the numbers of mosquitoes infected, with disseminated infections, infectiousness, and measures of transmission efficiency.

### The Effect of Viral Dose on ZIKV Transmission Efficiency

We measured the effect of viral dose on transmission efficiency; specifically, the proportion of infected mosquitoes that have disseminated infection (dissemination efficiency) and the proportion with disseminated infections that became infectious (efficiency of salivary gland infection). Despite the significant interaction between dose and day after infection on the proportion of mosquitoes with viral dissemination and saliva infection, dose and day did not affect these two response variables (Table 1, Fig 3). This indicates the midgut may be the primary barrier to infection, and that the rate of dissemination and salivary gland infection is not dose-dependent.

**Fig 3.**
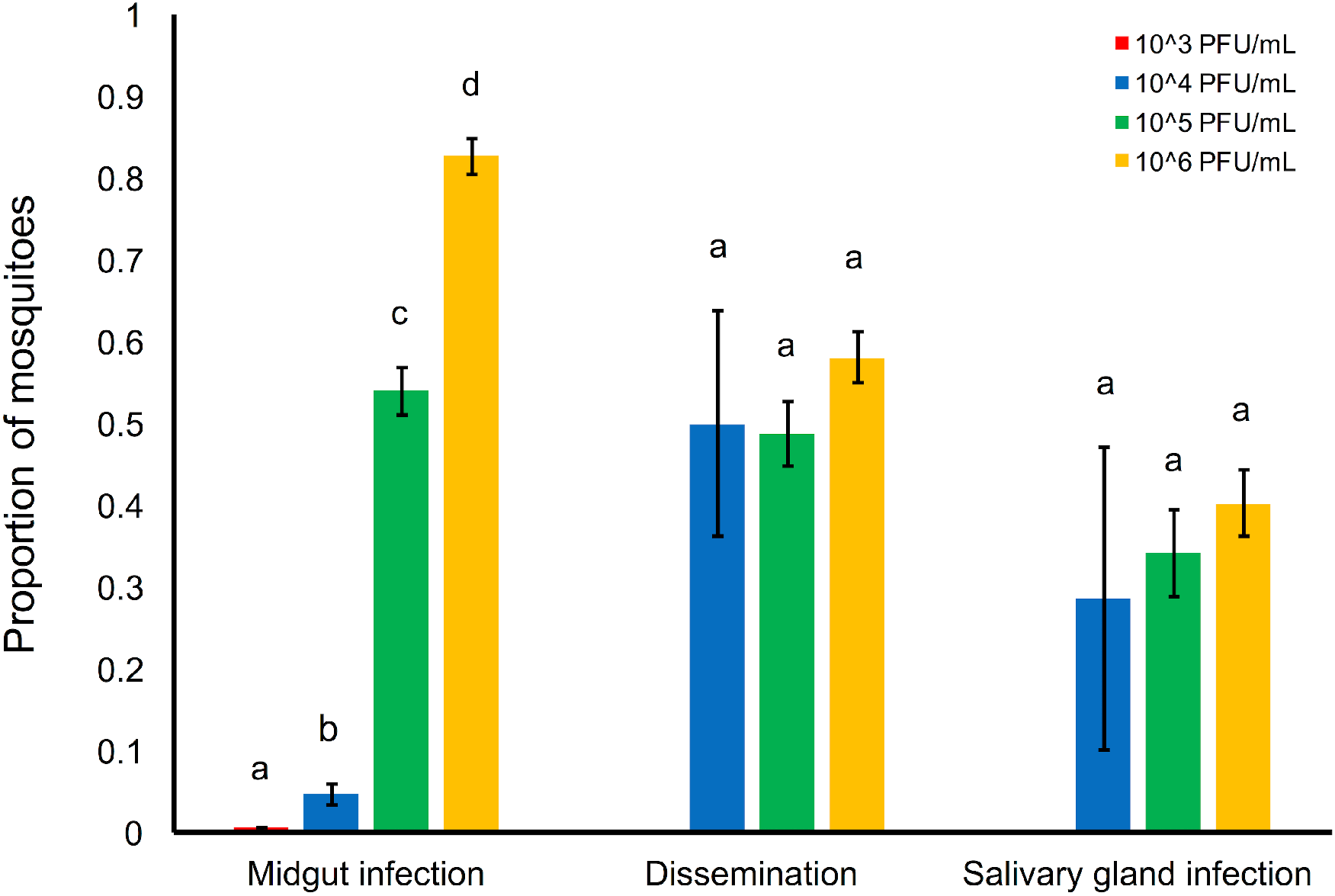
ZIKV dose and the efficiency of midgut infection, dissemination, and salivary gland infection. Relationship between the ZIKV dose (10^3^, 10^4^, 10^5^, and 10^6^ PFU/mL) and the efficiency of midgut infection (proportion of ZIKV positive bodies relative to the total number exposed), of dissemination (proportion of ZIKV positive heads relative to positive bodies), and salivary gland infection (proportion of ZIKV positive saliva compared to positive heads). For each category, results with no common letters were significantly different (*p* ≤ 0.05).

### The Effect of Zika Infection on Mosquito Survival

To determine if ZIKV infection and viral dose altered the daily probability of survival in *Ae. aegypti* mosquitoes, we included uninfected blood-fed controls in the study. We did not find any significant differences in the daily probability of survival between uninfected and ZIKV infected mosquitoes. Further, we observed no effects of increasing viral dose on mosquito survival among the infected mosquitoes (Table 2). On average, *Ae. aegypti* fed on viral doses of 10^3^, 10^4^, 10^5^, and 10^6^ PFU/mL experienced an average life span (*lf*) of 27, 24, 30, and 29 days, respectively.

**Table 2.**
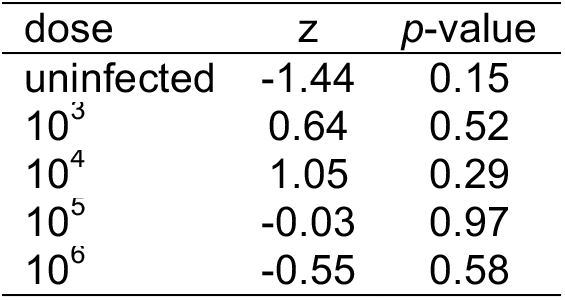
The effects of ZIKV dose on the daily probability of mosquito survival.

| dose | z | p-value |
| --- | --- | --- |
| uninfected | -1.44 | 0.15 |
| $10^3$ | 0.64 | 0.52 |
| $10^4$ | 1.05 | 0.29 |
| $10^5$ | -0.03 | 0.97 |
| $10^6$ | -0.55 | 0.58 |

Results from Cox mixed-effects model examining the effects of ZIKV dose (10^3^, 10^4^, 10^5^, and 10^6^ PFU/mL) on the daily probability of mosquito survival.

### Comparison between Plaque Assays and RT-qPCR

Most studies utilize qPCR to assess mosquito infection status. This method not only detects infectious particles, but also detects the viral genomic RNA in the infected cells, producing high RNA values that do not reflect the levels of infectious particles in the sample. When comparing the performance of plaque assays and RT-qPCR to assess infection status, we included the two highest doses (10^5^ and 10^6^ PFU/mL) because we had few to no positive saliva samples from the 10^3^ and 10^4^ treatment groups. Overall, the two methods gave similar numbers of positive samples (Table 3); however, we can detect the presence of ZIKV genome in mosquito saliva using RT-qPCR methods as early as 4 dpi, which was never the case with plaque assays.

In fact, infectious particles were rarely detected at 8 dpi with plaque assays. The number of infectious particles ranged from 3 (the limit of detection) to 120 PFU per sample and RNA molecules ranged from 10^4^ to 10^7^ gRNA copies (Fig 4A and Fig 4B). Both detection methods show that viral concentration does not have a significant effect on ZIKV levels in the saliva. However, we do see a significant effect of dpi on viral gRNA copies detected by RT-qPCR (Table 4).

**Table 3.**
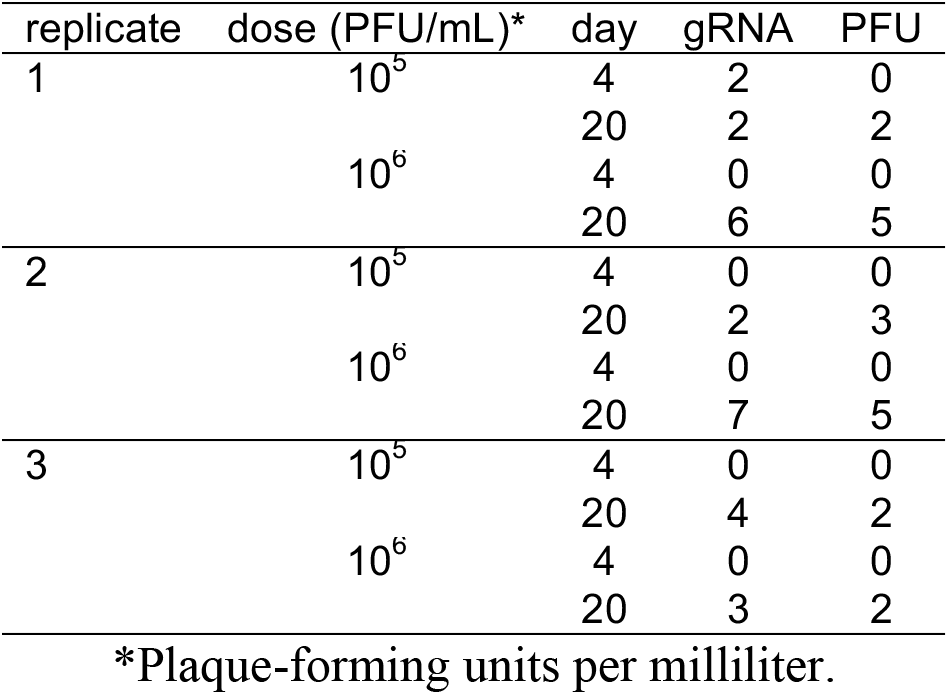
Numbers of positive saliva samples determined by RT-qPCR and plaque assays

**Fig 4.**
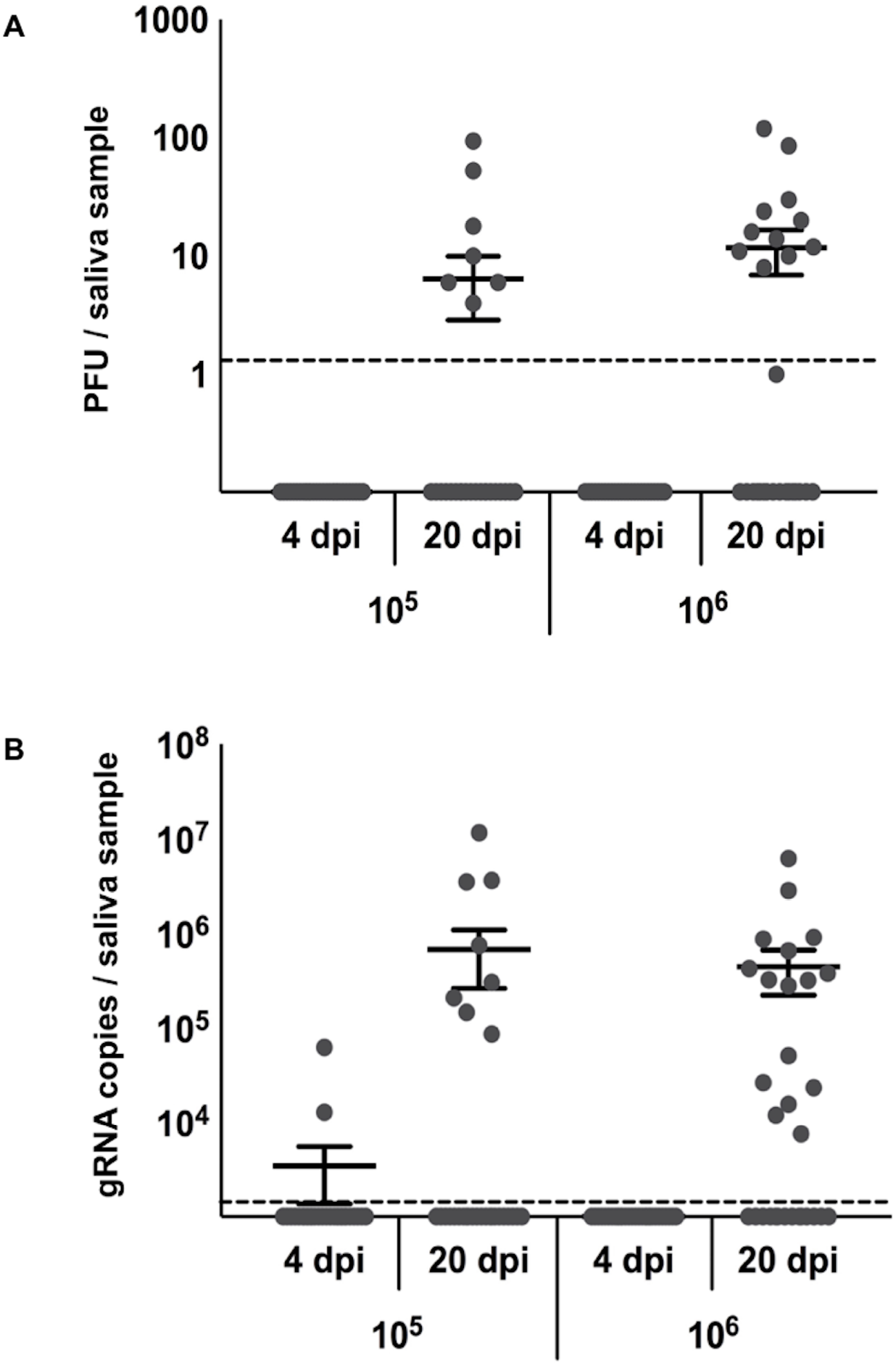
Viral loads in saliva determined by plaque assays and RT-qPCR. Viral load of ZIKV in saliva at 4 and 20 days post-infection (dpi) with 10^5^ and 10^6^ PFU/mL determined by standard plaque assays on Vero cells (A) and ZIKV-specific RT-qPCR (B). The limit of detection was experimentally established to be 3 plaque-forming units (PFU) for plaque assays and 30 gRNA copies for RT-qPCR.

Numbers of positive saliva samples determined by RT-qPCR (gRNA) and plaque assays (PFU) for 10^5^ and 10^6^ viral doses on days 4 and 20 post-infection for each experimental replicate.

**Table 4.** The effects of dose and day on the number of ZIKV gRNA copies and plaque-forming units

| factors | gRNA copies |  |  | plaque-forming units |  |  |
| --- | --- | --- | --- | --- | --- | --- |
|  | F | d.f. | p-value | F | d.f. | p-value |
| dose | 2.749 | 1 | 0.111 | 0.249 | 1 | 0.624 |
| day | 4.585 | 1 | **0.043** | - | - | - |

Results from generalized linear mixed effects models examining the effects of dose and day on the number of ZIKV gRNA copies vs. plaque-forming units

### The Effect of Viral Dose on Overall Transmission

The maximum proportion of the mosquito population that became infectious (vector competence; bc) increased with viral dose (Fig 5A). In contrast, the estimated *EIR* did not differ dramatically among mosquitoes fed different viral doses (Fig 5B), further suggesting that variation in infection dynamics with viral dose is driven primarily by positive dose effects on viral infection and escape from the midgut. This in turn resulted in increases in the relative transmission risk (*R_0_*) of mosquito populations feeding on hosts of increasing viremias (Fig 5C) and the relative number of infectious bites a human population would experience from a mosquito population of a given size (Fig 6).

**Fig 5.**
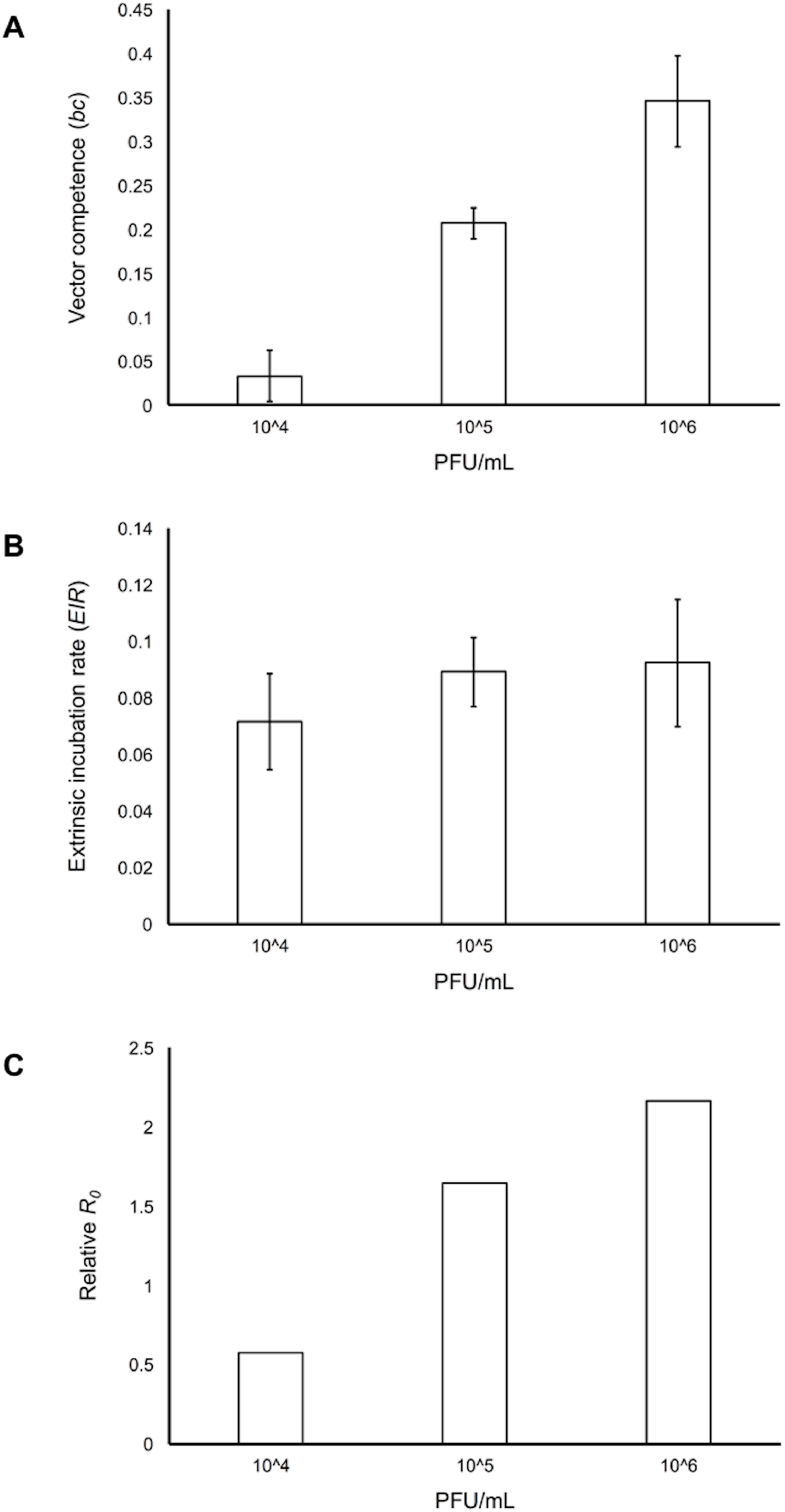
Viral dose and estimated vector competence, extrinsic incubation rate, and relative basic reproductive number *R_0_*. Relationship between viral dose (10^3^, 10^4^, 10^5^, and 10^6^ PFU/mL) and estimated vector competence (A), the extrinsic incubation rate (B), and relative basic reproductive number *R_0_* (C).

**Fig 6.**
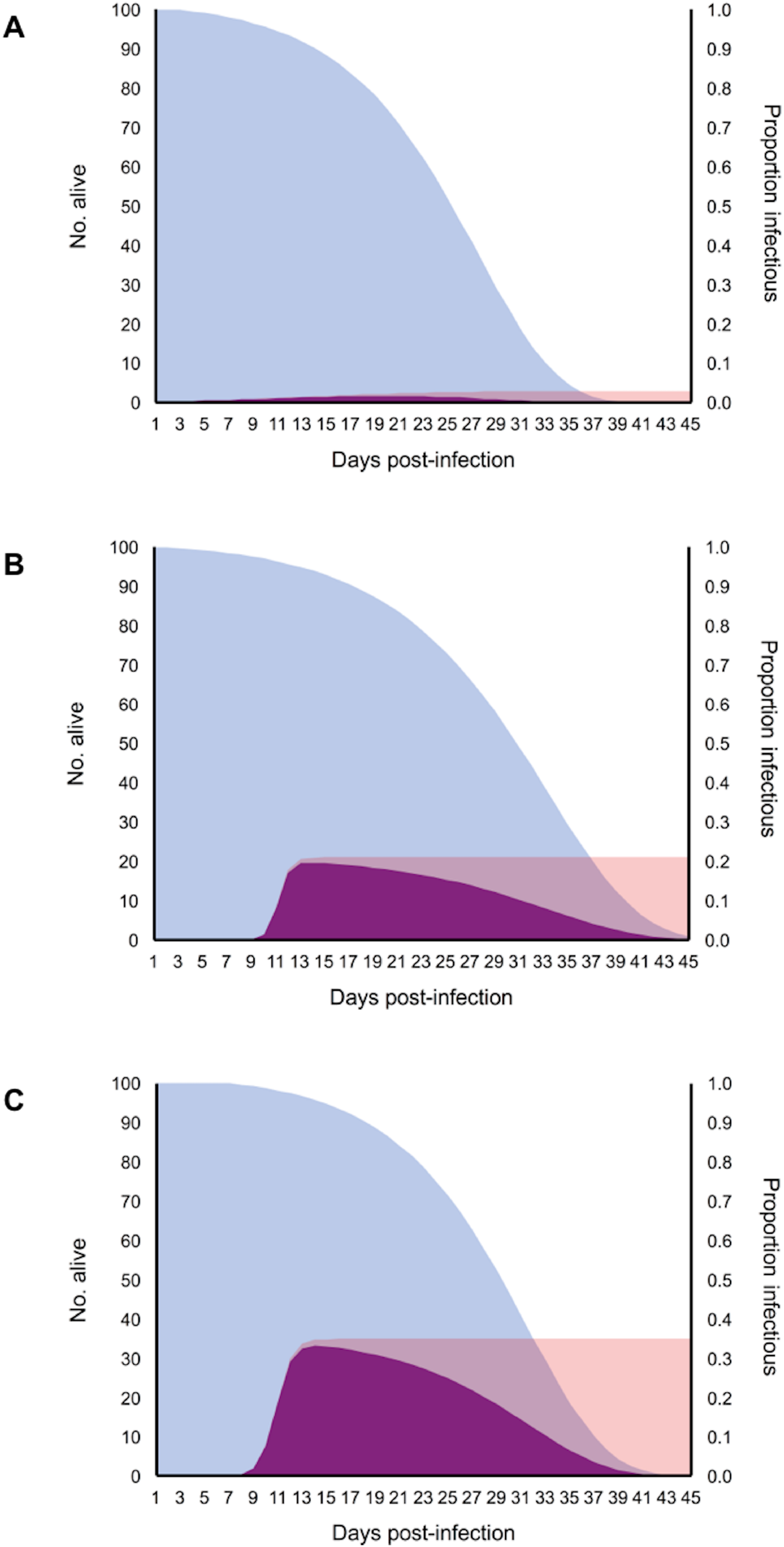
Daily proportion of mosquitoes alive, infectious, and both alive and infectious for mosquitoes exposed to different doses. Relationship between the daily proportion of mosquitoes alive (blue distributions), infectious (pink distributions), and those that are both alive and infectious (purple distributions) for mosquito populations exposed to 10^4^ PFU/mL (A), 10^5^ PFU/mL (B) and 10^6^ PFU/mL (C).

## Discussion

Mosquito vectors are often exposed to hosts that individually vary in pathogen loads, which can result in variation in the proportion of the mosquito population that becomes infectious (12, 26). To date, few studies have explored how viral concentration impacts measures of vector competence for ZIKV (13–15), and no studies have explicitly linked this source of variation to transmission risk. In this study, we demonstrate that *Ae. aegypti* populations exposed to increasing ZIKV concentrations exhibit increases in vector competence and EIR, which in turn results in substantial increases in relative transmission risk, measured as either *R_0_* or the force of infection.

Consistent with previous studies (13–15), we show that increasing the blood-meal concentration of ZIKV increases the probability mosquitoes will become infected, disseminate infection, and become infectious. Vector competence of a mosquito is strongly affected by the ability of a particular arbovirus to infect and escape the midgut and salivary gland barriers (27). As in other studies, we demonstrate that increases in viral concentration facilitates ZIKV infection and midgut barrier escape (28). A dose of at least 10^4^ PFU/mL was required for dissemination, and higher concentrations resulted in a higher proportion of mosquitoes with disseminated infections at earlier time points. Furthermore, we show that increases in viral concentration increases the EIR of ZIKV, consistent with other studies (15, 29). However, our study indicates that the effects of dose on the EIR is mainly through its positive effects on the ability of ZIKV to escape the midgut.

Compared to previous studies of ZIKV infection in *Ae. aegypti*, we found higher infection rates and a lower ID_50_. Our estimated ID_50_ (10^4.98^ PFU/mL) is much lower than previously reported ID_50_ 10^74^ PFU/mL (15). There is a substantial evidence in ZIKV, dengue, and chikungunya systems that vector competence can vary across mosquito populations due to genotype-by-genotype (G x G) interactions (14, 15, 29-31). Our higher infection rates could be due to the fact that we paired a Mexican ZIKV isolate with an *Ae. aegypti* population collected from the same region. Considering most ZIKV infected patients exhibit low viremia relative to other arboviruses, the mosquito-virus pairing may also explain why our infectious dose is more consistent with real-world viremias than previous estimates (14, 15, 32).

Mosquito longevity, along with EIR, are the strongest drivers of *R_0_*. Together these two parameters determine the duration of time a mosquito is alive and infectious. We found no effect of ZIKV infection or viral concentration on mosquito survival. It is generally assumed that mosquitoes are fairly tolerant of viral pathogens, allowing the virus to persist in the host without incurring fitness costs (33). However, most studies, including ours, have been performed in laboratory settings under relatively optimal conditions. Thus, if the costs of infection on mosquito survival and reproduction reflect underlying physiological trade-offs, fitness effects may only manifest in studies that incorporate relevant environmental stressors (e.g. variation in environmental temperature, food availability, competition, etc.) (34, 35).

To understand how variation in viral dose affects transmission risk, we used our infection and mortality data to parameterize a mechanistic *R_0_* model and determine the force of infection. Both *R_0_* and the force of infection are important measures of disease spread, representing the number of secondary cases in a susceptible population and the rate at which susceptible individuals acquire infectious disease, respectively. In our study, we show that mosquito populations feeding on increasing viral doses contribute more infectious bites and produce more secondary ZIKV cases due to increased vector competence and the rate at which virus escaped the midgut. For example, increasing viremia from 10^4^ to 10^6^ PFU/mL increased relative *R_0_* 3.8-fold and the number of infectious bites 18-fold. Knowing transmission risk will vary with heterogeneity in host viremia, future studies should focus on characterizing the distribution of viremia in the host population and incorporating individual variation in infectiousness into mechanistic models of disease spread. Model predictions from some pathogen systems (e.g. SARS, measles, and smallpox) that account for individual variation in infectiousness differ greatly from predictions generated by average-based approaches (36).

We used plaque assays to determine infection status instead of RT-qPCR, a common technique used in other studies due to its rapidity and sensitivity (37). However, because this method will detect all viral RNA in infected tissues, it can overestimate the actual number of infectious particles present. While we found both RT-qPCR and plaque assays to be highly correlated, the number of genomes detected by RT-qPCR was much higher than the number of plaque-forming units. We detected ZIKV genome in mosquito saliva (4 dpi) well before our first ZIKV infectious saliva sample was detected by plaque assay (8 dpi). Other studies using RT-qPCR methods have reported ZIKV in mosquito saliva as early as 3 dpi (13, 38). Since virus can be transmitted only in the form of infectious particles, the use of RT-qPCR to determine transmission relevant phenotypes could lead to overestimates of transmission risk.

In general, ZIKV viremia does not differ between symptomatic and asymptomatic patients (11) and is on average lower than seen with other arbovirus systems (29, 39). Contrary to our study, in the dengue and malaria systems, asymptomatic and pre-symptomatic patients with lower pathogen loads can be more infectious to mosquitoes than symptomatic hosts with high pathogen loads (12, 40). This could be due to host factors that are absent in our study and related studies (13, 15, 29). Variation in host blood quality (e.g. hematocrit) and mosquito attraction, or circulating host factors (e.g. differences in immune factors), could result in reduced infectivity of mosquitos feeding on hosts with high pathogen burdens (12, 41). Even the current, most frequently used ZIKV mouse models use mice lacking a large component of the innate immune system and are not likely to be representative of transmission in the field (14, 42). Thus, our study and others should be confirmed with mosquito feeding trials on human hosts of varying viremias.

In conclusion, we demonstrate that ingesting higher doses of ZIKV increases the proportion and the rate at which mosquito populations become infectious. This, in turn, results in an increase in the relative transmission risk and the force of infection experienced by susceptible human populations. Therefore, variation in viremia, as well as the frequency distribution of hosts of different viremias, should be accounted for when estimating *R_0_* and in assessing the efficacy of arbovirus prevention strategies.

## Materials and Methods

### Mosquito Rearing

We generated an outbred field-derived population of *Ae. aegypti* mosquitoes from ovitrap collections in Tapachula, Chiapas, Mexico, 2016. Larvae were reared in trays (200 larvae/1L ddH2O) and fed with 4 fish food pellets (Hikari Cichlid Cod Fish pellets). Larvae and adults were kept under standard insectary conditions at 27°C ± 0.5°C, 80%± 10% relative humidity, and a 12:12 hours light:dark diurnal cycle. Mosquitoes were maintained on human blood (Interstate Blood Bank) and provided with 10% sucrose *ad libitum*. F2 - F4 generations of mosquitoes were used for all downstream experiments.

### Virus Culture

For all mosquito infections, we used the ZIKV MEX1-44 strain obtained from the University of Texas Medical Branch Arbovirus Reference Collection. The virus was isolated from *Ae. aegypti* in 2016 from Chiapas, Mexico and passaged in Vero cells nine times. Vero cells were maintained in Dulbecco’s modified Eagle’s medium (DMEM) supplemented with 5% fetal bovine serum (FBS) at 37°C and 5% CO_2_. The virus was harvested four days after inoculation and stored at −80°C for at least seven days before titrating. Titers were determined by standard plaque assays on Vero cells as previously described (43), and expressed in plaque-forming units per milliliter (PFU/mL). Virus tested negative for *Mycoplasma* contamination using MycoSensor PCR Assay Kit (Agilent).

### Experimental Mosquito Infections

All ZIKV infections were performed under ACL3 conditions at the University of Georgia, Athens, GA, USA. Two days prior to infectious feed, mosquitoes were provided only water. On the day of infection, we prepared infectious and control blood-meals by washing human blood three times in RPMI medium. We then mixed 50% red blood cells with 33% DMEM, 20% FBS, 1% (wt/vol) sucrose, and ATP to a final concentration of 5 mmol/L. The blood-mixture was then mixed with virus at a 1:1 ratio. Three to five day-old mosquitoes were fed on a membrane feeder containing uninfected or infectious blood-meals with a final concentration of 10^3^, 10^4^, 10^5^ or 10^6^ PFU/mL for 30 min. We then randomly distributed 40 engorged mosquitoes across five 16 oz paper cups for each dose. The mosquitoes were fed 10% sucrose *ad libitum* for the duration of the experiment, and we recorded the number of dead mosquitoes every two days. Three biological replicates were performed.

### Quantifying Mosquito Infection via Forced Salivations

To determine the proportion of mosquitoes that were infected with ZIKV, had disseminated infections, and were infectious, we processed 20 mosquitoes per treatment group on days 4, 8, 12, 16 and 20 post-infection. Mosquitoes were cold anesthetized and kept on ice until their legs and wings were removed. After immobilization, mosquitoes were transferred to a hot plate (35°C) where they salivated into a tip containing 35 μL FBS with 3 mmol/L ATP and red food dye for 45 min. After salivation, mosquitoes were decapitated, and bodies, heads, and saliva were individually placed into tubes containing 600 μL of DMEM with 1x antibiotic/antimycotic. Bodies and heads were homogenized in a QIAGEN TissueLyzer at 30 cycles/second for 30 seconds, and centrifuged at 17,000xg for 5 minutes at 4°C. Samples were then assessed for the presence/absence of virus with plaque assays.

### RT-qPCR Analysis

To compare plaque assays with quantitative reverse transcription PCR (RT-qPCR), we performed RT-qPCR on ten saliva samples per replicate at days 4 and 20 post-infection from mosquitoes exposed to 10^5^ and 10^6^ PFU/mL. Viral RNA was extracted from saliva samples (QIAamp^®^ Viral RNA Mini Kit, Qiagen) and reverse-transcribed to cDNA (High Capacity RNA-to-cDNA Kit, Applied Biosystems). ZIKV genome copies were measured with RT-qPCR reaction assay using TaqMan^®^ Gene Expression Master Mix (Applied Biosystems), primers (F: ZIKV 1086, R: ZIKV 1162c; Invitrogen Custom Primers) and probes (ZIKV 1107-FAM; TaqMan MGB Probe) (44). Each sample was analyzed in duplicate, and each assay contained a standard curve (ZIKV molecular clone), no template, and no primer controls. We extrapolated ZIKV copy numbers from the generated standard curve using the Applied Biosystems protocol. The limit of detection was experimentally established to be 30 copies (10^−16^ g). Final copy numbers were adjusted by back-calculations to the total RNA and cDNA volume and expressed as copies per saliva sample.

### Statistical Analysis

We used mixed effects generalized linear models (IBM^®^ SPSS^®^ Statistics 1.0.0.407) to estimate the effects of ZIKV dose, day post-infection (dpi), and the interaction (fixed factors) on the number of mosquitoes becoming infected (positive bodies: negative binomial distribution, log link function), disseminating infection (positive heads: normal distribution, identify link function), and becoming infectious (positive saliva: normal distribution, identity link function). Similar models were also constructed to assess dose and dpi effects on transmission efficiency (of those infected, the number with disseminated infections, and of those with disseminated infections, the number with positive saliva: Poisson distribution, log link function) and viral burdens in the saliva (normal distribution, log link function). Finally, we used a Cox mixed effects model (R version 3.3.3, package ‘coxme’ (45)) to estimate the effects of ZIKV infection, dose, and the interaction on the daily probability of mosquito survival. Experimental replicate was included in all models as a random factor. Model fit and distributions were determined based on Akaike Information Criterion (AIC), the dispersion parameter, and by plotting model residuals. Sequential Bonferroni tests were used to assess the significance of pairwise comparisons when relevant, and p-values greater than 0.05 were considered non-significant.

### Mechanistic *R_0_* Model

To estimate the effects of dose on transmission risk, we used two different approaches. First, we calculated relative *R_0_* as a function of dose (*x*) since the absolute magnitude of *R_0_* depends on other factors not considered here. We modified a function of *R_0_* used in previous work (17) (Equation 1):

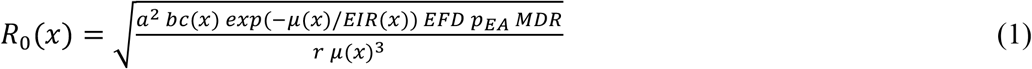

with parameters for the daily biting rate (*a*), vector competence (*bc*), the daily adult mosquito mortality rate (*μ*), the extrinsic incubation rate (*EIR*), the probability of egg to adult survival (*p_EA_*), the mosquito development rate (*MDR*), and the human recovery rate (*r*). For parameters we did not directly estimate (*a*, *p_EA_*, *MDR*), we used estimates generated by Mordecai et al. (17) for *Ae. aegypti* at 27°C and assumed the human recovery rate to be the inverse of the average number of days ZIKV is detectable in the blood (6 days (46)). We used the experimental infection data to estimate dose-dependent vector competence (bc), the *EIR*, and the daily mortality rate (*μ*) as follows. We fit logistic growth models to the proportion of infectious mosquitoes (*Y*) versus dpi (*t*) for each viral dose (*x*) (Equation 2),

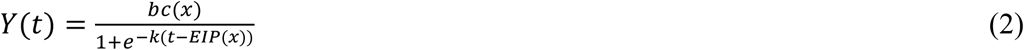

using the “nls” package in R (47). Vector competence was defined as the maximum proportion of infectious mosquitoes achieved per dose (the asymptote, *bc*), *EIR* was estimated as the inverse of the extrinsic incubation period (the inflection point, *EIP*), and *k* reflects the instantaneous rate of increase (slope at the inflection point). Then, to estimate the daily probability of mosquito mortality (*μ*) we fit a variety of non-linear curves (exponential, log-linear, Weibull, and Gompertz) to the daily survival probabilities of mosquitoes exposed to different doses with the “flexsurv” package in R (48). We used AIC to determine the best performing model and calculated the area under the curve to estimate the average lifespan (*lf*) of mosquitoes exposed to varying doses. The average daily probability of mortality was then estimated as the inverse of the dose-specific lifespan (1/*lf*).

Second, we performed an alternative calculation of transmission risk following previously described methods (49) to account for the substantial variation in infection outcomes observed across mosquitoes exposed to a given dose. Briefly, we multiplied the best fitting nonlinear functions describing the daily relationship between survival and the proportion of infectious mosquitoes for each dose treatment, resulting in the number of infectious days/dose. We then estimated the area under the curve of the resulting function and multiplied by the daily biting rate (*a*) (17) to calculate the number of infectious mosquito bites generated for each dose treatment for a mosquito population of a given size (n=100).

## Acknowledgments

We thank the University of Texas Medical Branch Arbovirus Reference Collection for providing the virus. We also thank the members of the Murdock and Brindley labs for thoughtful comments on the project and manuscript. Any opinions, findings, and conclusions or recommendations expressed in this material are those of the author(s) and do not necessarily reflect the views of the National Science Foundation.

This study was supported by the National Science Foundation, Grants for Rapid Response Research (NSF-RAPID) 1640780. Erin A. Mordecai was supported by NSF DEB-1518681 and the Stanford University Woods Institute for the Environment Environmental Ventures Program.

